# “Lead (Pb) impairs thyroid hormone mediated changes in brain development and body length in *Xenopus laevis* tadpoles”

**DOI:** 10.1101/2022.09.27.509775

**Authors:** Lara I. Dahora, Alayna M. Robinson, Christopher Buenaventura, Hannah Bailey, Christopher K. Thompson

## Abstract

Lead (Pb) poisoning during early development is associated with behavioral and cognitive deficits, but the specific mechanisms by which Pb impairs brain development are not fully understood. One potential mechanism is that Pb poisoning may impair thyroid hormone (TH)- mediated changes in brain development To address this issue, we performed experiments to assess the effects of Pb poisoning on (TH) -dependent changes in cellular and molecular mechanisms in the developing *Xenopus laevis* tadpole brain. We treated stage 48 tadpoles to combinations of 1000 ppb Pb bath for seven days and added one of three different concentrations of thyroxine (T_4_) for the final two days of treatment. We found that lead exposure decreased body length, including in T_4_-treated tadpoles. We also performed immuno-staining for proliferative marker pH3 and found that Pb disrupts T_4_-induced increases in neuronal proliferation. Finally, we used syGlass VR data visualization software to measure volume of the forebrain, midbrain, and hindbrain in 3D and found that Pb exposure impaired T_4_-mediated changes in brain volume. Last, we found that Pb poisoning reduced the T_4_-mediated increase in proliferating cell nuclear antigen (PCNA), a TH-sensitive gene. These results illustrate that Pb poisoning impairs some TH-dependent changes in the developing brain.

## INTRODUCTION

One of the most pernicious effects of lead (Pb) poisoning in children is compromised brain development, leading to behavioral impairments with major societal costs. Children with blood lead levels 5-10μg/dL score on average 4.9 points lower on Full-Scale IQ (Jusko *et al*., 2008). Further, there is a significant dose-dependent association between childhood blood lead levels and the risk of a learning disability diagnosis (Geier *et al*., 2017). The Health Impact Project estimates that prioritizing the elimination of environmental lead poisoning in children could alleviate $84 billion in economic burden in the form of decreased healthcare costs and financial harm to affected families (Health Impact Project, 2017). The CDC estimates children in at least 4 million households are exposed to high levels of Pb through drinking water, and the Department of Housing and Urban Development estimates 37 million residences contain Pb-contaminated paint (Centers for Disease Control and Prevention, 2017; US Dept. of HUD, 2018).

Despite these findings, we still lack a complete picture of exactly how Pb poisoning affects the developing brain. Several studies on the effects of Pb on the developing central nervous system (CNS) have highlighted an increase in cell death. For instance, Pb poisoning induces the loss of rod and bipolar cells in the retina as a result of apoptosis during development (Fox *et al*., 1997). This study found that the retina is acutely sensitive to lead exposure, observing effects both in developing and adult animals. Pb also increases lipid peroxidation in the striatum, thalamus, and hippocampus of developing rats, which could also lead to cell death (Villeda-Hernández *et al*., 2001). Because Pb^2+^ substitutes for Ca^2+^, Pb may stimulate calmodulin, leading to inhibition of neurite development (Kern and Audesirk, 1995). Not much else is known about the effects of Pb on other cellular attributes in the developing CNS.

While there are significant gaps in our understanding of the effects of Pb poisoning on the developing CNS, we do know that acute exposure to Pb impairs adult brain function in multiple ways. Pb exposure alters expression of synaptic proteins and electrophysiological properties (Neal *et al*., 2010; Neal and Guilarte, 2013; Xiao *et al*., 2006; Yamada *et al*., 1995; Omelchenko *et al*., 1996). In addition, Pb exposure affects calcium-dependent processes and the expression of neurotrophic factors (Lasley and Gilbert, 1996, 2002). Some of these deficits could have the potential to impair the developing CNS.

While it is clear that Pb poisoning directly impedes many critical molecular pathways that underlie brain development, the effects of Pb poisoning on thyroid hormone (TH) physiology in the developing brain are not well understood. Pb can decrease circulating levels of thyroxine (T_4_), triiodothyronine (T_3_) thyroid stimulating hormone (TSH) in children and in adults (Dundar *et al*., 2006; Abdelouahab *et al*., 2008). More importantly, elevated blood lead levels are associated with lower levels of circulating T_4_ in pregnant women (Kahn *et al*., 2014). The developing CNS requires adequate levels of TH, and lower levels of TH pre- and postnatally are associated with cognitive dysfunction and behavioral disorders, not unlike what has been observed in Pb-exposed children. Another study found that lead-induced hypothyroidism and impairment of long-term potentiation in hippocampal CA1 region of developmental rats is rescued with levothyroxine, a synthetic version of T_4_ (Wu *et al*., 2011). Their results showed that Pb exposure led to a decrease in T_3_, T_4_, and a dramatic decrease in TSH and administering levothyroxine restored these to control levels. This may be particularly important to the developing brain because TH is a regulator of a variety of critical developmental milestones in the brain (Patel *et al*., 2011). Exposure to Pb has various effects on adult thyroid hormone physiology. Pb can decrease levels of thyroid hormones and differentially regulate thyroid hormone-related genes (Miao *et al*., 2015).

The goals of this study were to evaluate the effects of Pb on thyroid hormone-mediated mechanisms of brain development. Specifically, we assessed the effect Pb had on neuronal proliferation, brain volume, body length, and TH-sensitive gene expression. Our results show that Pb impaired TH-mediated increases in neuronal proliferation and body length. Additionally, Pb abolished TH-mediated increase in PCNA gene expression in the brain. Last, Pb affected TH-mediated changes in brain volume. This suggests that Pb can inhibit thyroid hormone-mediated changes in the brain.

## METHODS

### Animals

We used albino *Xenopus laevis* tadpoles bred on site. Hatching was carried out in 20-gallon tanks maintained at 18°C under a 12:12-hour light:dark photoperiod. We fed tadpoles once per day beginning when yolk was depleted (~7 dpf). We used stage 48 tadpoles (10-12d postfertilization [dpf]), which were selected based on morphological characteristics in accordance with Nieuwkoop and Faber’s *Xenopus* table of development (1956, 1994). Once selected, tadpoles were placed in glass bowls (200 mL each) in Steinberg’s rearing solution. During the course of the experiment, we kept the tadpoles in a 22°C incubator and maintained the 12:12-h light:dark cycle.

All animal procedures were conducted in accordance with Virginia Tech’s Institutional Animal Care and Use Committee’s rules and regulations.

### Experimental Treatments

To systemically treat tadpoles with lead (Pb), 500 mg of lead (II) acetate trihydrate (Sigma: 215902-25G) was diluted into 50 mL of hot water to create a stock solution of 26.36 mM or 10,000,000 parts per billion (ppb) lead. Stock solution was stored at room temperature until used in experiments. Tadpoles were placed into glass bowls (200 mL each) in Steinberg’s rearing solution. To bring the concentration of lead (Pb) in experimental groups to 1000 ppb, 20 μL of the stock solution was added to 200 mL Steinberg’s for the duration of the seven-day experiments.

To systemically treat tadpoles with the thyroid hormone thyroxine (T_4_), we dissolved 100 μg of crystalline T_4_ (Sigma) in 6.66 mL of 50 mM NaOH (1.93 mM) and further diluted it to 1.93 μM in Steinberg’s solution to make a stock solution, which was stored at −20°C. Aliquots of 1.93 μM T_4_ stock were thawed, and 20 μL, 200 μL, and 2 mL were added to 200 mL (0.15 μg/L and 1.5 μg/L concentrations) and 198 mL (15 μg/L concentration), to bring bowls to final T_4_ concentrations of 0.15 μg/L, 1.5 μg/L, and 15 μg/L (19.3 nM), respectively. T_4_ was added to bowls on the fifth day of experimental treatment with lead (Pb) for thyroid hormone treatment on the final two days of the experiments.

### Euthanasia and Fixation

Tadpoles were euthanized by 0.2% of tricaine methanesulfonate (MS-222; Sigma) overdose, then fixed in 4% phosphate-buffered paraformaldehyde (PFA) at least overnight at 4°C before dissection. Those utilized in quantitative real-time PCR were euthanized by 0.2% MS-222 as well, prior to dissection of the brain.

### Body Length

To quantify body length, we imaged fixed animals in 4% PFA and imaged them on a Nikon SMZ 745T dissecting microscope using a USB 3.0 color CMOS 2.3 MP camera. Then, we measured the body lengths of fixed tadpoles from snout to tail tip in ImageJ (RRID: SCR_003070).

### Whole-Mount Immunohistochemistry, Tissue Clearing, and Imaging

Following euthanasia and fixation, we stained dissected brains with antibodies to assess neurogenesis as well as with Vybrant™ DiD cell-labeling solution (Invitrogen; V2289) to visualize overall brain volume. After fixation in 4% PFA, we dissected out brains and placed them into phosphate buffered saline with 0.1% Triton-X (PBS-0.1% TX). We then did two washes with PBS-0.1% TX for fifteen minutes each. Following the initial washes, we permeabilized the brains for one hour in PBS-2.0% TX-100. We then performed another wash in PBS-0.1% TX-100. Next, we blocked non-specific binding using 2.5% normal goat serum in PBS-0.1% TX-100 for one hour with gentle shaking. We then incubated brains in primary anti-phosphorylated histone 3 (pH3) (Sigma-Aldrich; H0412) antibody solution (in PBS-0.1% TX-100) at a concentration of 1:1000 overnight at 4°C. The following day, we performed three fifteen-minute washes in PBS-0.1% TX-100. After washes, we incubated brains in secondary antibody solution (Invitrogen AlexaFluor™ 488 goat anti-rabbit; A11034) at a concentration of 1:400 for four hours with gentle shaking. After secondary antibody incubation, we performed two more fifteen-minute washes prior to incubating brains in 1:1000 concentration DiD in DMSO for three days in order to visualize the whole brain and assess total brain volume.

We cleared dissected brains using the X-FaCT (Xenopus-Fast Clearing Technique) method (Affaticati *et al*., 2018). Briefly, this is a technique where we immersed brains in a fructose-based high-refractive index solution (fbHRI) in order to clear the tissue and use confocal imaging to get higher quality 3D images of whole brains.

Brains were then mounted in wells on a slide for imaging on a Leica SP8 confocal microscope. An emission based Z-correction was applied and the images are montages reassembled by the Leica LASX software package (RRID: SCR_013673).

### Quantitative real-time PCR

To quantify the changes in the expression of thyroid hormone (TH)-sensitive genes, we placed stage 48 tadpoles in a 2 x 4 experimental paradigm bath. In one dimension, tadpoles were treated with either 1000 ppb lead (Pb) or control bath (Steinberg’s solution); in the other dimension, tadpoles were placed in 0 μg/L (CNTL), 0.15 μg/L, 1.5 μg/L, or 15 μg/L thyroxine (T_4_). This experimental design provides a control group and a 1000 ppb lead (Pb) group for each concentration of T_4_. After 7 days of treatment with lead (Pb), T_4_ of varying concentrations was added to bowls for the final two days of the experiment. After 4d, we euthanized the tadpoles by overdose of MS-222 (0.2%); quickly dissected out the brains and placed them into Trizol (Life Technologies), with 3 brains per tube; and froze them at −80°C until day of RNA extraction. We extracted RNA in accordance with the protocol provided by the manufacturer of Trizol and performed RNA cleanup when necessary. Following extraction, RNA quality and concentrations were measured on a NanoDrop One (Thermo Scientific) for all samples. Samples of RNA were then reverse-transcribed u sing the Bio-Rad iScript kit with 500 ng of RNA per reaction. We then performed quantitative polymerase chain reaction (qPCR) using 2 ng of complementary DNA for each reaction using the iTaq Universal SYBR Green Supermix kit (Bio-Rad) on a Bio-Rad CFX384 thermocycler. Primers used for quantification of NREP, PCNA, and klf9 can be found in Table 1. We used a 2-step reaction with 10s 95°C melt step, followed by a 30s 60°C annealing and an extension step for 40 cycles, with fluorescence measured at the end of every 60°C step.

**TABLE 1.**
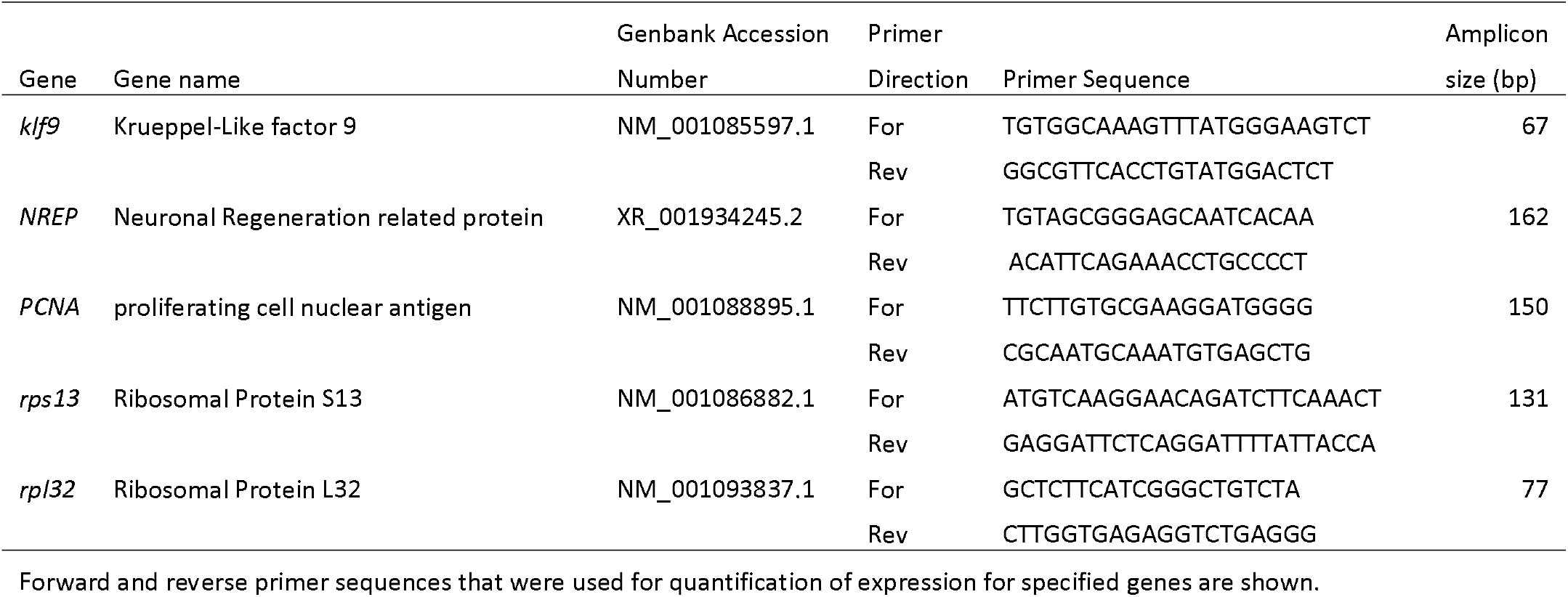
Primers used for quantification of gene expression.

Data from qPCR experiments were taken from Bio-Rad CFX software following runs and moved to Microsoft Excel for analysis. All reactions were done in triplicate, and outliers with deviations >1.5 times the SD from the mean within a set of triplicates were removed from analysis. The reference genes used were rps13 and rpl32; the primers used for these genes are also included in Table 1. These ribosomal proteins were selected as reference genes for normalization because their expression was stable and showed no statistical differences between groups. They have been validated as stable reference genes for *Xenopus* tadpoles in a previous study (Thompson and Cline, 2016). The mean levels of expression for each gene were normalized to the controls for their respective backgrounds and are reported as a fold change relative to these groups. They were also normalized to the reference genes by subtracting the average of both genes for each sample. Expression of target genes was evaluated using the ΔΔCq method against the mean expression per sample of these genes.

### SyGlass VR Analysis (Volume and Proliferation)

Quantification of proliferating cells (pH3+ cells) were done using SyGlass Data Visualization software (RRID: SCR_017961). To quantify number of proliferating cells, we used the counting tool in SyGlass to count each individual cell in the forebrain, tectum, and hindbrain separately. In order to quantify volume of each area, we used the carving tool and carved out the volume of those three areas of the brain. In order to quantify whole brain volume and number of proliferating cells, we added all of them together to get the measure of the full brain.

### Quantification and Statistical Analysis

Statistical analyses were made using GraphPad Prism, Ver 8.4.3 (GraphPad Software, RRID: SCR_002798). Data sets with more than two groups were evaluated with analysis of variance (ANOVA), and post hoc comparisons were made with Tukey’s multiple comparisons test. Data sets with two dimensions (e.g., lead treatment vs thyroxine treatment) were evaluated with □ way ANOVA, and post hoc comparisons were made with Tukey’s multiple comparisons test, Sidak’s multiple comparisons test, or Dunnett’s test when multiple groups were compared with a control group when appropriate.

## RESULTS

### Effects of Pb Exposure on Body Length with Exogenous Administration of Thyroxine

To evaluate the effects of Pb exposure on body length, we exposed stage 48 tadpoles to control (Steinberg’s rearing solution) or 1000ppb Pb for seven days. Additionally, we added three different concentrations of T_4_: 0.15 μg/L, 1.5 μg/L, or 15 μg/L for the final two days of the experiment. The control group was unexposed to Pb or T_4_ the entire time course. Generally, administration of TH induces an initial increase in body length followed by a decrease in body length as tail absorption proceeds (Thompson and Cline, 2016). Pb-exposed animals in all groups exhibited shorter body length than their non-exposed counterparts (Figure 1A). Furthermore, this decrease in body length in Pb-treated groups appeared to be dose-dependent in terms of the amount of exogenous T_4_ administered the final two days of the experiment. The longest average body length is in the T_4_ 15μg/L group that was not treated with Pb, while the shortest body length seen is in the T_4_ 15μg/L that was treated with 1000ppb Pb. Representative images of tadpoles from each experimental group are shown in Figure 1B. There were statistically significant differences between all CNTL and Pb-treated groups with exogenous administration of 0 μg/L, 0.15 μg/L, 1.5 μg/L, or 15 μg/L, with p= 0.0012, 0.0001, <0.0001, and <0.0001, respectively (2way ANOVA, Sidak’s multiple comparisons test).

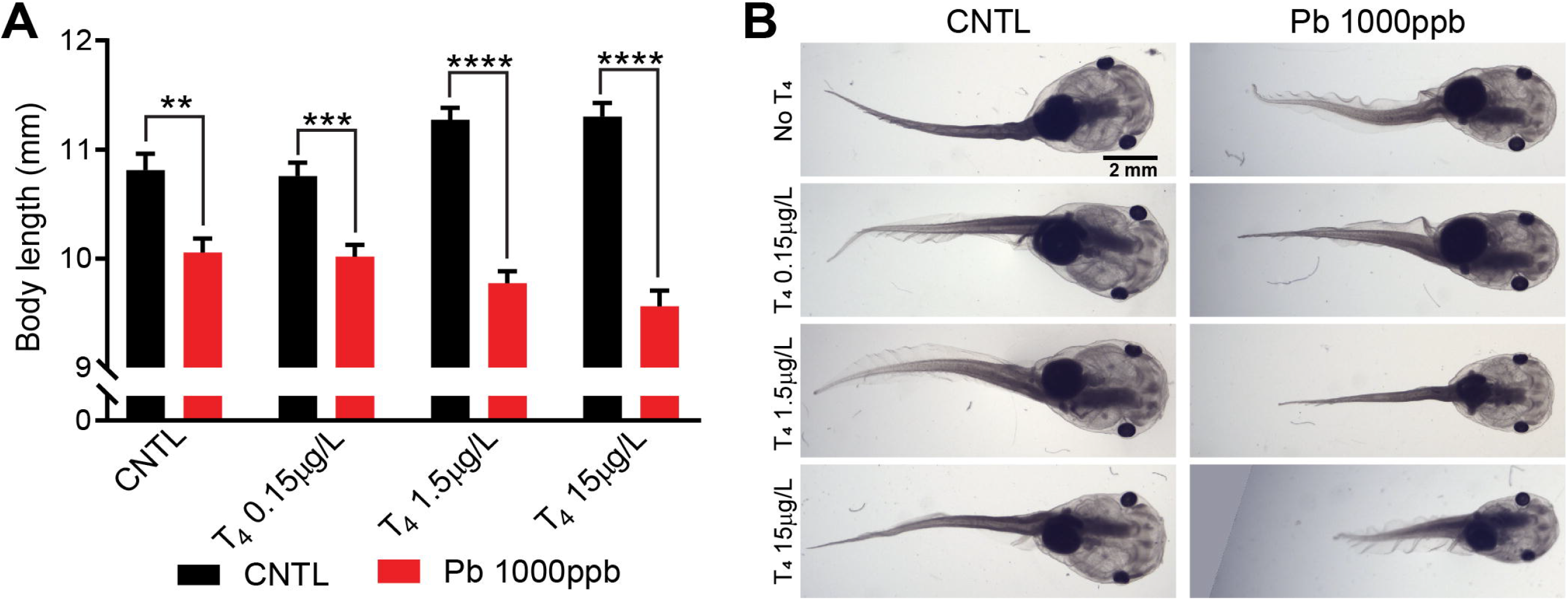

### Effects of Pb Exposure with Exogenous Administration of Thyroxine on Thyroid Hormone-Sensitive Gene Expression

To evaluate the effects of Pb exposure on thyroid hormone dependent gene expression, we performed qPCR to quantify expression of proliferating cell nuclear antigen (PCNA), neuronal regeneration-related protein (NREP), and Krueppel-like factor 9 (klf9) in the brains of animals treated with either control bath or Pb (1000ppb) for seven days. Additionally, we exogenously administered the same three concentrations of T_4_: 0.15 μg/L, 1.5 μg/L, or 15 μg/L for the final two days of the experiment. No statistically significant differences were seen in expression of NREP between lead-treated and non-treated animals in any group, regardless of T_4_ treatment. Marginally significant differences were seen in klf9 at lower concentrations of T_4_. Statistically significant differences were seen in the T_4_ 1.5 μg/L and T_4_ 15μg/L groups between CNTL and lead-treated groups for PCNA, with p= 0.0037 and 0.0001, respectively (2way ANOVA, Sidak’s multiple comparisons test).

### Effects of Pb Exposure with Exogenous Administration of Thyroxine on Cell Proliferation in the Developing Brain

Administration of T_4_ induces a dose-dependent increase in proliferation in all areas of the brain (divided here into forebrain, tectum, and hindbrain) (Thompson and Cline, 2016). To evaluate the effect of Pb exposure on neuronal proliferation during brain development, we exposed stage 48 tadpoles to control or Pb (1000ppb) for seven days. Additionally, we exogenously administered three different concentrations of T_4_: 0.15 μg/L, 1.5 μg/L, or 15 μg/L for the final two days of the experiment. We found the lowest number of pH3+ cells in the control group not treated with any T_4_, and the highest number of pH3+ cells in the non-Pb-treated T_4_ 15μg/L group (Figure 3B). The difference between non-Pb-treated groups and Pb-treated groups in forebrain proliferation was greatest in the T_4_ 15μg/L groups. Specifically, there were minor differences in number of pH3+ cells in the T_4_ 0μg/L and T_4_ 0.15μg/L groups, but the difference between CNTL- and 1000ppb + T_4_ 15μg/L group was statistically significant (p= 0.001). Quantification of pH3+ cells in the tectum revealed a substantial decrease in proliferation between non-Pb- and Pb-treated groups that had either T_4_ 1.5 μg/L or T_4_ 15μg/L added the final two days of the experiment. The difference between CNTL and 1000ppb Pb in the T_4_ 1.5 μg/L and T_4_ 15μg/L groups was statistically significant with p= 0.0279 and <0.0001, respectively (Figure 3C). The largest differences between non-Pb-treated and Pb-treated groups was seen in the hindbrain. Drastic differences were revealed in the groups treated with 1.5 and 15μg/L T_4_ for the final two days of the experiment, with statistically significant differences for both groups at p < 0.0001 (Figure 3D). Finally, we combined data from the three brain regions in order to ascertain changes in whole brain proliferation. Pb-treatment almost completely suppressed T_4_-stimulated proliferation in the 1.5 or and 15μg/L T_4_-treatment groups. The differences between those treated and not treated with Pb were statistically significant with a p < 0.0001 (2-way ANOVA, Sidak’s multiple comparisons test). These data show clearly that Pb treatment almost completely abolishes T_4_-induced increases in proliferation when compared with non Pb-treated animals (Figure 3E).

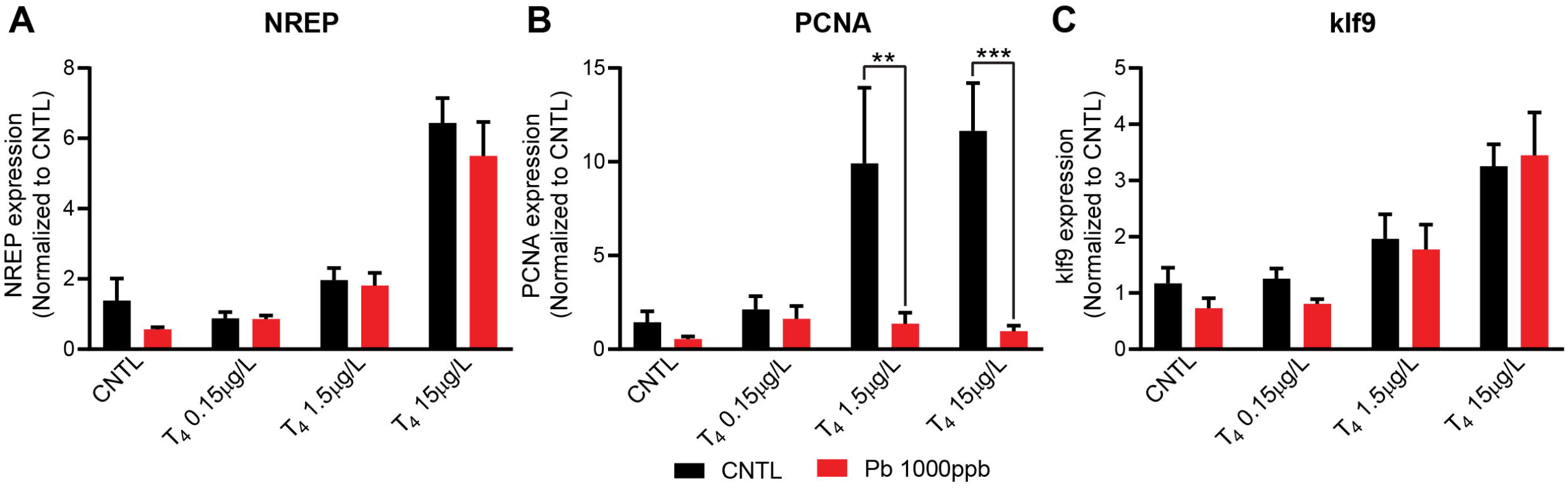

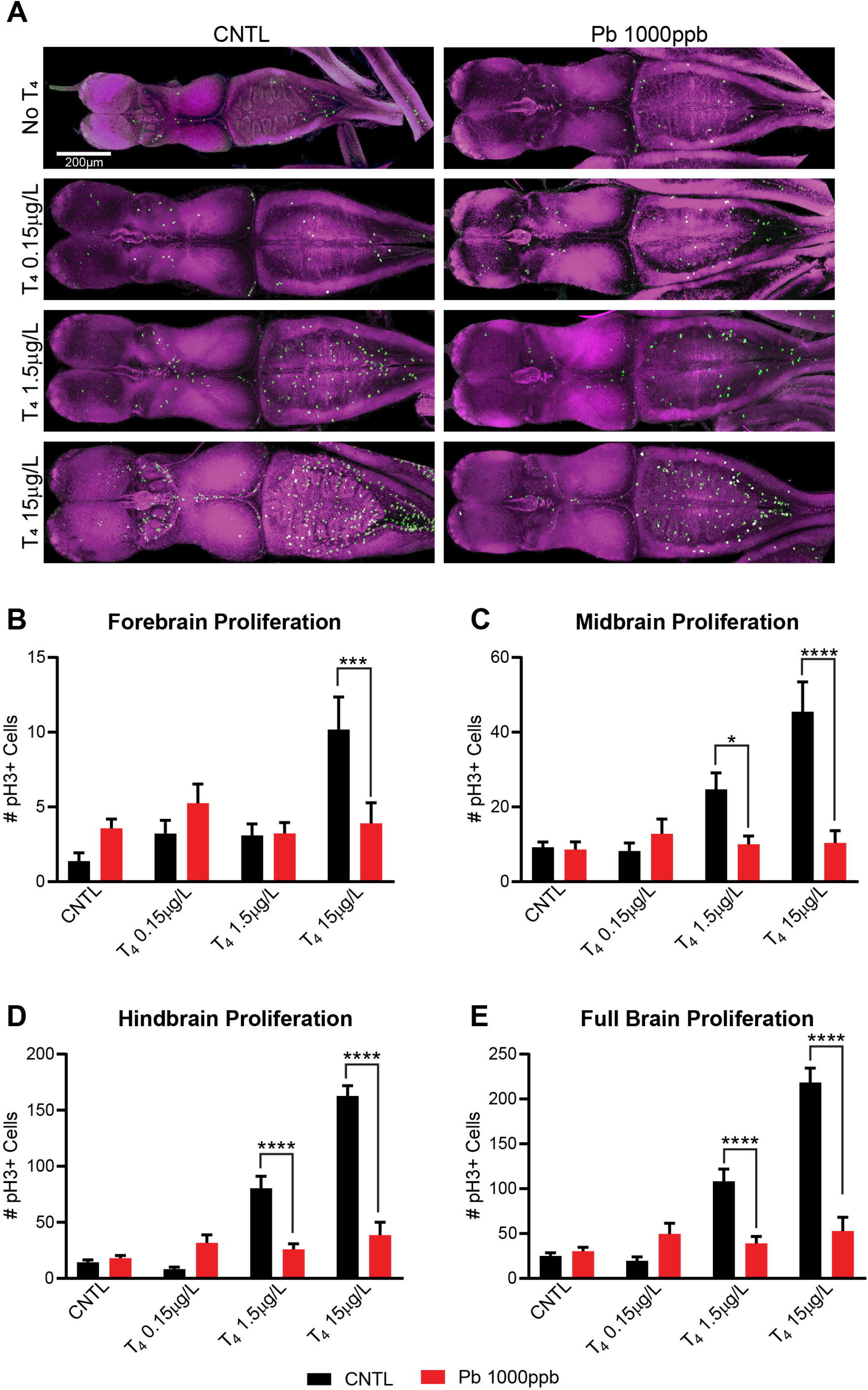

### Effects of Pb Exposure with Exogenous Administration of Thyroxine on Brain Volume During Development

To evaluate the effect of Pb exposure on brain volume during brain development, we followed the same paradigm used in the proliferation experiment but imaged those same brains with DID staining in order to quantify changes in brain volume, using the same images that were used for quantifying proliferation. Example images depicting volume renderings, including proliferation quantification points, are shown in Figure 4A. Volume of CNTL animals not treated with T_4_ was lower than those treated with Pb (1000ppb) in all areas of the brain including forebrain (p= 0.497; 2way ANOVA, Sidak’s multiple comparisons test), tectum (p=0.2475), and hindbrain (p=0.1001), although the difference in the forebrain was the only brain region that showed statistical significance. Measured volume in the CNTL group treated with 0.15μg/L was also lower than the 1000 ppb Pb-treated group that was also treated with 0.15μg/L, although the difference was not statistically significant (p=0.9579). There were statistically significant differences in forebrain volume between both the T_4_ 1.5 and 15μg/L groups treated with CNTL and 1000 ppb Pb bath, with p= 0.0003 and 0.0167, respectively.

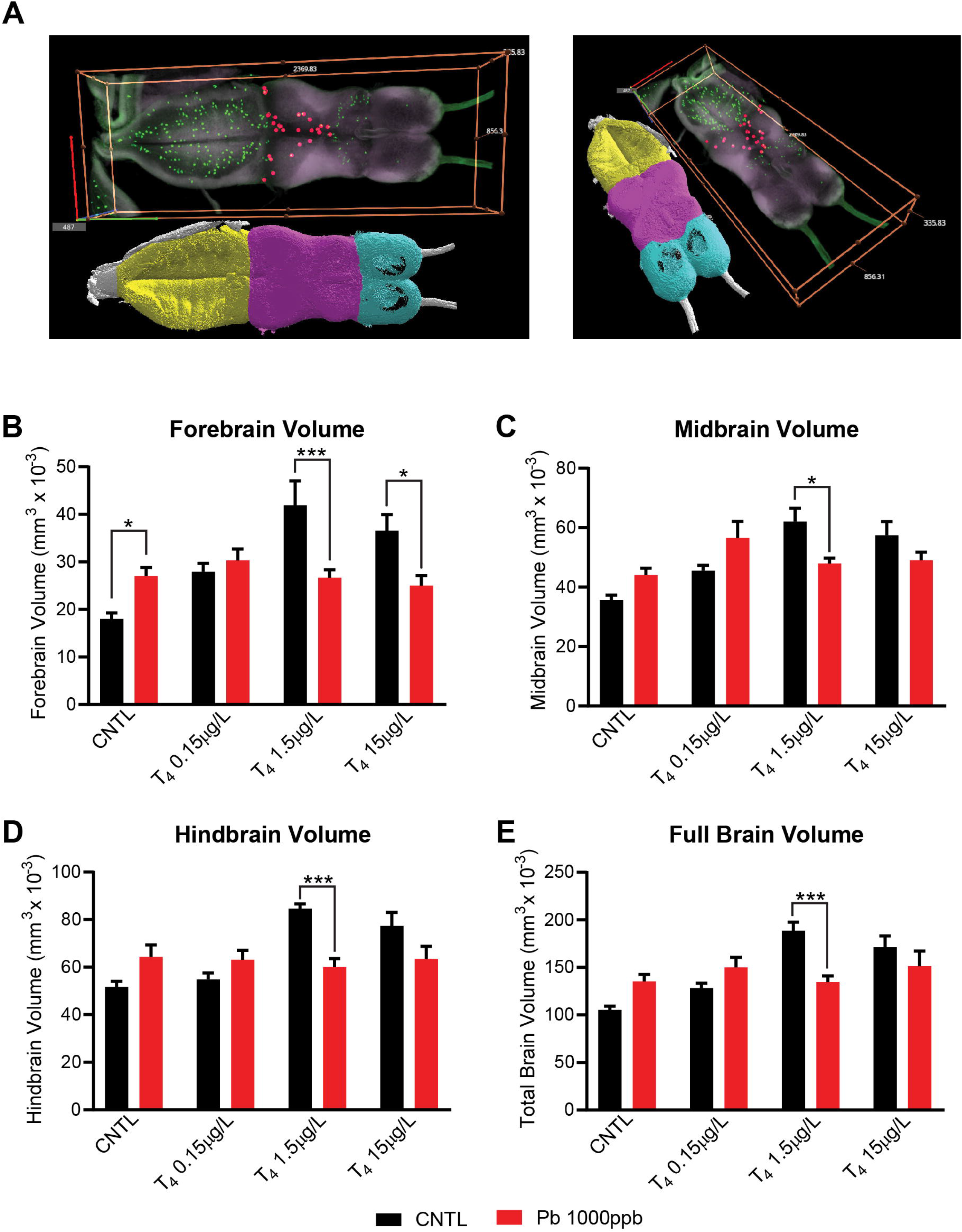

## DISCUSSION/CONCLUSIONS

Our results show that Pb treatment impaired TH-induced changes in neuronal proliferation and changes in brain volume in *X. laevis* tadpoles. Results also show Pb-treatment impaired body length. Furthermore, Pb decreased in PCNA expression, consistent with the decrease seen in neuronal proliferation. Finally, our results show no statistically significant differences between CNTL and Pb-treated groups in klf9 or NREP expression. Collectively, these data show impairment of thyroid hormone-induced increases in neurogenesis and brain volume in response to systemic Pb exposure.

The decrease in brain volume, although not uniform, may be indicative of a change in overall brain anatomy than just changes in volume. One study found that in adults that were exposed to lead as children had decreased brain volume due to a loss in gray matter (Cecil *et al*., 2008). Their findings were based on MRI and changes were seen in prefrontal cortex and anterior cingulate cortex, including areas responsible for mood regulation, executive function, and decision-making. Another MRI study found that past adult lead (Pb) exposure was associated with decreased volumes in the cingulate gyrus and insula (Stewart *et al*., 2006). While the general trend of the data showed a decrease in brain volume in response to Pb treatment, we observed an increase in the 1000ppb Pb treated group in forebrain volume relative to untreated tadpoles. This could be because there is limited proliferation in the forebrain compared to the mid- and hindbrain, and there could have been inflammation in that group that the T_4_-treated groups overcame. Previous work has shown that lead exposure induces inflammation in rat pups, which could account for this increase in brain volume that we observed in the CNTL 1000ppb Pb group (Chibowska *et al*., 2020).

Our data clearly shows that Pb treatment significantly impaired T_4_-induced neuronal proliferation, suggesting that Pb likely impedes mechanisms that regulate proliferation in the brain. Specifically, the decrease is most apparent in the two groups with the highest T_4_ concentrations. While forebrain proliferation was decreased only in the highest T_4_ concentration group, midbrain and hindbrain proliferation, as well as full brain proliferation was seen to decrease in the two highest T_4_-treated groups. These two groups were exposed to supernormal concentrations of T_4_ at these stages and normally induce a drastic increase in proliferation in response to thyroid hormone treatment. These data are in line with previous research that has shown the capacity for Pb to inhibit neurogenesis in both young animals and adult animals. Gilbert et al. showed that chronic low-level Pb exposure reduced neurogenesis in the adult hippocampus (Gilbert *et al*., 2005). Additionally, early life exposure to Pb significantly decreased neurogenesis in the adult brain (Verina *et al*., 2007), and Pb treatment in embryonic zebrafish undergo impaired neurogenesis (Dou and Zhang, 2011). Likewise, another study found a decrease in the number of PCNA positive cells postnatally in animals treated with Pb compared to controls although Pb exposure significantly increased PCNA positivity during gestational days (Mousa *et al*., 2018). Nevertheless, most of the studies evaluating the effects of Pb on neurogenesis focused on the effects on adult neurogenesis in rat models. A study in juvenile rodents found that the number of synapses was significantly less than that of the control group in animals treated with Pb both before weaning and after weaning (Xiao *et al*., 2014). This same study also found changes in synaptic structural parameters such as thickness of postsynaptic density, length of synaptic active zone, synaptic curvature, and width of the synaptic cleft compared to the control group (Xiao *et al*., 2014). Not every study finds that Pb-exposure decreases neurogenesis, however. For instance, gestational Pb exposure increased retinal progenitor and bipolar cell proliferation in rodents, which may be reactive neurogenesis to account for increased cell death (Giddabasappa *et al*., 2011).

There are two conclusions to be drawn from the body length results. First, it is very clear that Pb-treatment impaired overall growth, including in the comparison that does not involve exposure to T_4_. The mechanism by which Pb toxicity impaired growth is less clear, however. A study in humans in Yugoslavia showed that low level childhood exposures to Pb were associated with decreased height (Factor-Litvak *et al*., 1999), similar to other studies that also found decreased height and body mass index (BMI) in response to Pb exposure (Deierlein *et al*., 2019; Scinicariello *et al*., 2013). One study, however, failed to find an association of blood lead levels and height but instead found decreases in offspring height and weight in mothers who had elevated thyroid hormone during pregnancy (Lamb *et al*., 2008). Furthermore, Pb poisoning is negatively associated with head circumference and stature (Ballew *et al*., 1999). Given that our results show that Pb-significantly impairs proliferation in the brain, Pb may similarly impair proliferation in other areas of the body. The second conclusion to be drawn is little more subtle. TH treatment induces changes in body length in tadpoles that are not unidirectional. There is first elongation and then, within a few days, shortening of body length upon precocious systemic treatment with thyroid hormone (Thompson and Cline, 2016). Given that the tadpoles in are experiment are exposed to T_4_ for two days, we observed a dose-dependent increase in body length as expected. Interestingly, body length in Pb-treated tadpoles in fact decreased with increasing concentrations of T_4_, in the opposite direction of T_4_ treatment alone. This suggests that Pb toxicity is exacerbated upon exposure to T_4_. The mechanism by which this is occurring is difficult to say, and future experiments will need to address this issue.

The specific effects of Pb on thyroid hormone physiology and signaling is not well understood, particularly during development. Several studies have examined the effects of Pb on overall thyroid hormone physiology, mostly in adults, but the effects are not clear as there have been a wide range of results, some of which are conflicting when attempting to draw a clear picture of the effects. Some of the studies show hypothyroid-like levels of thyroid hormones in response to Pb exposure, while others show hyperthyroid-like levels of thyroid hormones or no effects on thyroid hormones at all (Chen *et al*., 2013; Yousif and Ahmed, 2009; Fahim *et al*., 2020; Sahin *et al*., 2016; Erfurth *et al*., 2001; Wu *et al*., 2011). Because of the lack of a consensus on the effects of Pb on thyroid hormone physiology, it remains to be seen what the actual effects of Pb are during development across species.

There are various potential mechanisms of neurotoxicity when thinking about the effects of Pb exposure. As previously mentioned, Pb exposure affects NMDAR subunit expression, leading to changes in electrophysiological properties (Yamada *et al*., 1995; Omelchenko *et al*., 1996). Another potential mechanism of neurotoxicity is the effect that Pb has on thyroid hormone physiology in the brain itself. Our data suggest that thyroid hormone distribution in the brain of *Xenopus laevis* tadpoles may be at least partially impaired by Pb toxicity. It is possible that this impairment is due to decreased thyroid hormone making its way to the brain due to decreases in thyroid hormone distributor proteins. Previous work in rats has shown that chronic, low-dose Pb exposure is associated with significant decreases in cerebrospinal fluid (CSF) thyroid hormone distributor protein transthyretin (TTR), which is produced by the choroid plexus (Zheng *et al*., 1999, 2001). Additionally, the effects of Pb on thyroid hormone physiology may compromise energetics in proliferating cells. Previous work has already demonstrated effects of Pb on mitochondrial function and morphology (Zhang *et al*., 2009). Zhang and colleagues demonstrated that Pb exposure led to mitochondrial swelling, cristae loss, and disruption. Further, it has been shown that Pb has biphasic effects on mitochondrial respiration, with low levels of exposure leading to an increase in respiration and higher concentrations of Pb exposure inhibiting this lead-induced increase in respiration (Holtzman *et al*., 1978). It is possible that these effects of Pb on mitochondrial function and morphology could be effected through thyroid hormone-mediated mechanisms. Other mechanisms of neurotoxicity include decreases in calcium-dependent glutamate and GABA release, decreases in expression of presynaptic synaptophysin and synaptobrevin, decreases calcium influx through voltage-gated calcium channels, and decreases in BDNF expression (Lasley and Gilbert, 1996, 2002; Xiao *et al*., 2006; Neal *et al*., 2010; Neal and Guilarte, 2013; Stansfield *et al*., 2012).

Overall, our results show that Pb poisoning impairs many aspects of how thyroid hormone acts in the developing tadpole, including changes in brain morphology, proliferation, gene expression and over body length. Our data suggest that Pb may have the ability to impair thyroid hormone-mediated changes in brain development in other animals. Future work will need to assess the mechanisms by which these changes are effected and will need to evaluate whether these impairments are due to changes in thyroid hormone distributor proteins or other changes in thyroid hormone physiology.

## FUNDING FOR THIS WORK

This work is funded by NIEHS grant R00ES022992. This work is also funded by NIEHS fellowship F31ES031855.

